# Resistance to Ceftazidime/Avibactam Plus Meropenem/Vaborbactam When Both are Used Together Achieved in Four Steps From Metallo-β-Lactamase Negative *Klebsiella pneumoniae*

**DOI:** 10.1101/2020.03.09.983304

**Authors:** Punyawee Dulyayangkul, Edward J. A. Douglas, Filip Lastovka, Matthew B. Avison

## Abstract

Serine cephalosporinases and carbapenemases are dominant causes of critically important β-lactam resistance in *Klebsiella pneumoniae*. This has led to the recent clinical deployment of new serine β-lactamase inhibitors used in combination with β-lactams. Starting with clinical *K. pneumoniae* isolates and adding plasmids carrying the OXA-48-like class D carbapenemase, OXA-232, the class A carbapenemase KPC-3, the class A cephalosporinase CTX-M-14 and mutant derivatives of these enzymes, we set out to identify the steps required to give resistance to the recently approved β-lactam/β-lactamase inhibitor pairs ceftazidime/avibactam and meropenem/vaborbactam when both are used together. We show that four steps: *ompK36* and *ramR* loss-of-function plus carriage of OXA-232 and KPC-3-D178Y, all of which have been observed in clinical isolates, allow *K. pneumoniae* to resist the combined use of both β-lactam/β-lactamase inhibitor pairs. These findings have implications for decision making about sequential and combinatorial use of β-lactam/β-lactamase inhibitor pairs to treat *K. pneumoniae* infections, and suggest simple surveillance activities that might identify intermediate stages in resistance acquisition and therefore guide therapy to reduce the emergence of dual resistant strains.

## Introduction

Cephalosporin- and carbapenem-resistant *Klebsiella pneumoniae* are critically important pathogens (1). *K. pneumoniae* can become resistant to a variety of antimicrobials by increased production of two RND efflux pumps, AcrAB and OqxAB (2–5) or reduced production of two porins, OmpK35 and OmpK36 (6). *K. pneumoniae* can also acquire a wide range of serine β-lactamases (7) the most clinically significant of which are enzymes of the CTX-M (class A cephalosporinase [8]), KPC (class A carbapenemase/cephalosporinase [9]) and the OXA-48-like (class D carbapenemase [10]) types.

In response to the rise of CTX-M and KPC, serine β-lactamase inhibitors have been developed. Avibactam contains a diazabicyclo-octane cyclic core (11) and is used in combination with the third-generation cephalosporin ceftazidime. In vitro, avibactam shows good activity against many class A (TEM-1, SHV-1, CTX-M and KPC), class C (AmpC) and even some class D (OXA-24 and OXA-48) β-lactamases (12,13). When used *in vivo*, ceftazidime/avibactam resistance has been seen to develop in KPC producers, caused by a series of point mutations in the *bla_KPC_* gene, for example resulting in D179Y or V239G substitutions in KPC (14,15). A similar mechanism of ceftazidime/avibactam resistance was seen in CTX-M producers, with mutations in *bla*_CTX-M_, for example leading to a P170S change, alone or in addition to T264I (16).

Vaborbactam is a cyclic boronate and is used in combination with the carbapenem meropenem (17). It has potent activity against class A carbapenemases including KPC. It has also been shown to have inhibitory activity against other class A (TEM, SHV and CTX-M) and class C (AmpC) β-lactamases, although to a lesser extent when compared to its activity against KPC (18). Meropenem/vaborbactam resistance has been observed due to loss of function mutations in *ompK36* and *ompK35* (18,19). Mutations in KPC that confer reduced vaborbactam inhibition have not yet been observed.

Given the appearance of *K. pneumoniae* clinical isolates and laboratory selected mutants that are resistant to meropenem/vaborbactam or ceftazidime/avibactam, there has been some discussion in the literature about whether a combination of both β-lactam/β-lactamase inhibitor pairs given together would overcome isolates resistant to either, and to both when used separately (20,21). In the work reported here we identified the steps required to generate resistance to each β-lactam/β-lactamase inhibitor pair, both pairs when used separately, and both pairs when used together. To do this we started with *K. pneumoniae* clinical isolates susceptible to both meropenem and ceftazidime.

## Results and Discussion

### Permeability mutations influencing Meropenem/Vaborbactam MICs in K. pneumoniae Ecl8 carrying OXA-232 or KPC-3

There are four main loss-of-function mutations that affect accumulation of antibacterial drugs in *K. pneumoniae* clinical isolates: loss of the transcriptional repressors OqxR (2,3) and RamR (4,5), and loss of the porins OmpK35 (22,23) and OmpK36 (24,25). Starting with a laboratory workhorse *K. pneumoniae* strain, Ecl8, we determined the influence of these mutations on meropenem MIC in the presence or absence of 8 μg.mL^−1^ vaborbactam when the strains carried one of two carbapenemases commonly encountered in the clinic: the OXA-48-like enzyme OXA-232 (26) or the class A carbapenemase KPC-3 (15). OXA-232 is a relatively weak carbapenemase, and in our hands does not give meropenem resistance in this otherwise wild-type background (**Table 1**) as we saw in the same background when using OXA-48 (27). Each loss-of-function mutation conferred a 4-fold increase in meropenem MIC, but in no case was the mutant meropenem resistant. In the presence of vaborbactam, meropenem MIC only marginally reduced, which was expected because vaborbactam is a poor inhibitor of OXA-48-like enzymes (28). In contrast, KPC-3 is an excellent carbapenemase and gives meropenem resistance even in this wild-type background (**Table 1**). All four loss-of-function mutations increased meropenem MICs, but only by one or two doubling dilutions. Vaborbactam is a strong inhibitor of KPC-3 (17), and meropenem susceptibility was restored in Ecl8 and all mutants producing this enzyme. The highest MIC was against the *ompK36* mutant, but it remained meropenem/vaborbactam susceptible. This was expected, since it has been shown that meropenem/vaborbactam resistance in derivatives of *K. pneumoniae* clinical isolates is primarily caused by *ompK36* loss in a background where *ompK35* has already been lost (19) and we have previously reported that Ecl8 actually produces more OmpK35 than clinical isolates (29).

**Table 1.** MICs (μg.mL^−1^) of meropenem with or without vaborbactam against *K. pneumoniae* Ecl8 derivatives.

|  | **Meropenem** | **Meropenem/Vaborbactam** |
| --- | --- | --- |
| **Ecl8(pOXA-232)** | 0.125 | 0.125/8 |
| **Ecl8 *ompK35*(pOXA-232)** | 1 | 0.5/8 |
| **Ecl8 *ompK36*(pOXA-232)** | 1 | 1/8 |
| **Ecl8 *ramR*(pOXA-232)** | 1 | 0.5/8 |
| **Ecl8 *oqxR*(pOXA-232)** | 1 | 0.5/8 |
| **Ecl8(pKPC-3)** | 64 | 0.125/8 |
| **Ecl8 *ompK35*(pKPC-3)** | 128 | 0.125/8 |
| **Ecl8 *ompK36*(pKPC-3)** | 256 | 2/8 |
| **Ecl8 *ramR*(pKPC-3)** | 128 | 0.125/8 |
| **Ecl8 *oqxR*(pKPC-3)** | 128 | 0.125/8 |

Shading indicates resistance according to CLSI breakpoints (34).

### Generating derivatives of K. pneumoniae clinical isolates resistant to ceftazidime/avibactam or meropenem/vaborbactam but not both

*K. pneumoniae* clinical isolate KP21 produces low levels of OmpK35 (29). Experimentally, it has been shown that *ramR* loss of function downregulates OmpK35 production in *K. pneumoniae* (27) and KP21 carries a frameshift mutation in *ramR* (29) whilst whole genome sequencing confirmed that *ompK35* itself is wild type. We introduced a natural OXA-232 plasmid into this isolate, and the MIC of meropenem in the presence of vaborbactam against KP21(pOXA-232) was 1 μg.mL^−1^ (**Table 2**) which is below the resistance breakpoint and equivalent to our findings for Ecl8 *ramR*(pOXA-232) (**Table 1**). We next selected a spontaneous mutant of KP21(pOXA-232) which grew on 16 μg.mL^−1^ meropenem in the presence of vaborbactam, and so it is resistant. Proteomics confirmed that this mutant, KP21 M(pOXA-232), had undetectable OmpK36 porin levels, and whole genome sequencing identified a single mutation resulting in a stop at codon 125 in *ompK36*, explaining loss of the porin. The MICs of meropenem with or without vaborbactam against KP21 M[*ompK36*](pOXA-232) and against the constructed mutant KP21 *ompK36*(pOXA-232) were 256 μg.mL^−1^ (**Table 2**). This shows that *ompK36* loss confers meropenem/vaborbactam resistance in *ramR* mutant *K. pneumoniae* clinical isolates (i.e. with reduced OmpK35 levels) producing OXA-48-like enzymes. Loss-of-function mutations in *ompK35* are not necessary for this phenotype.

**Table 2.**
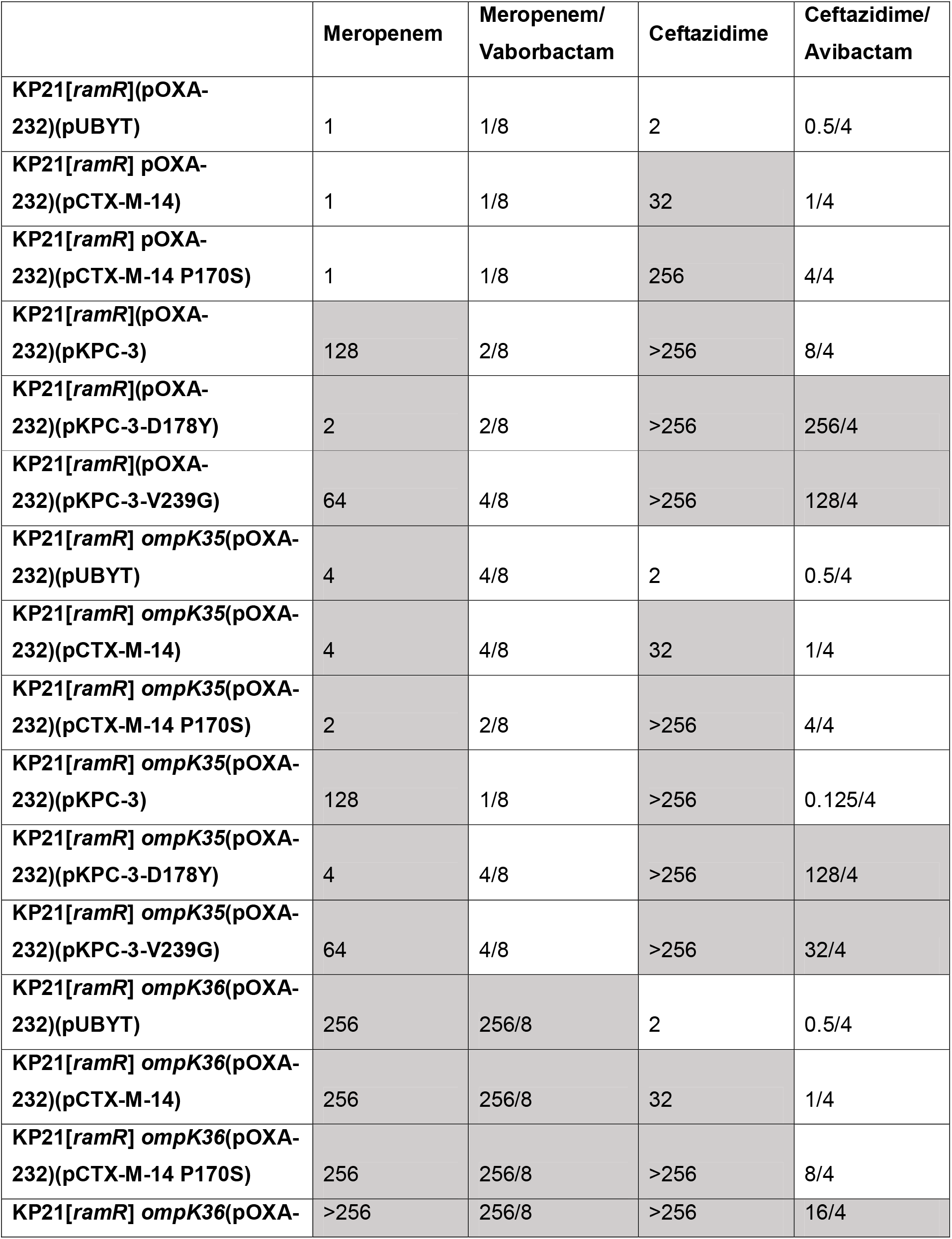
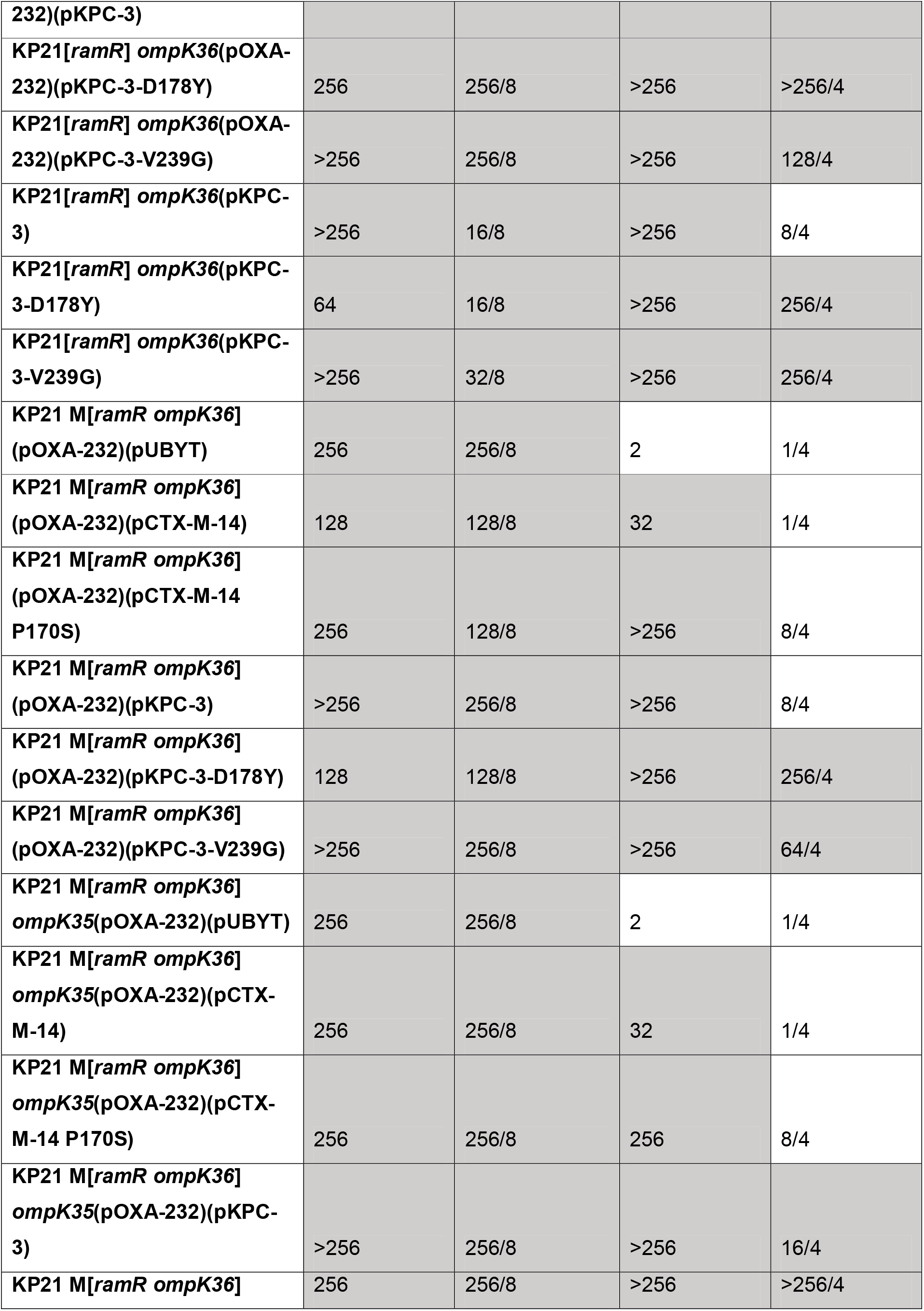
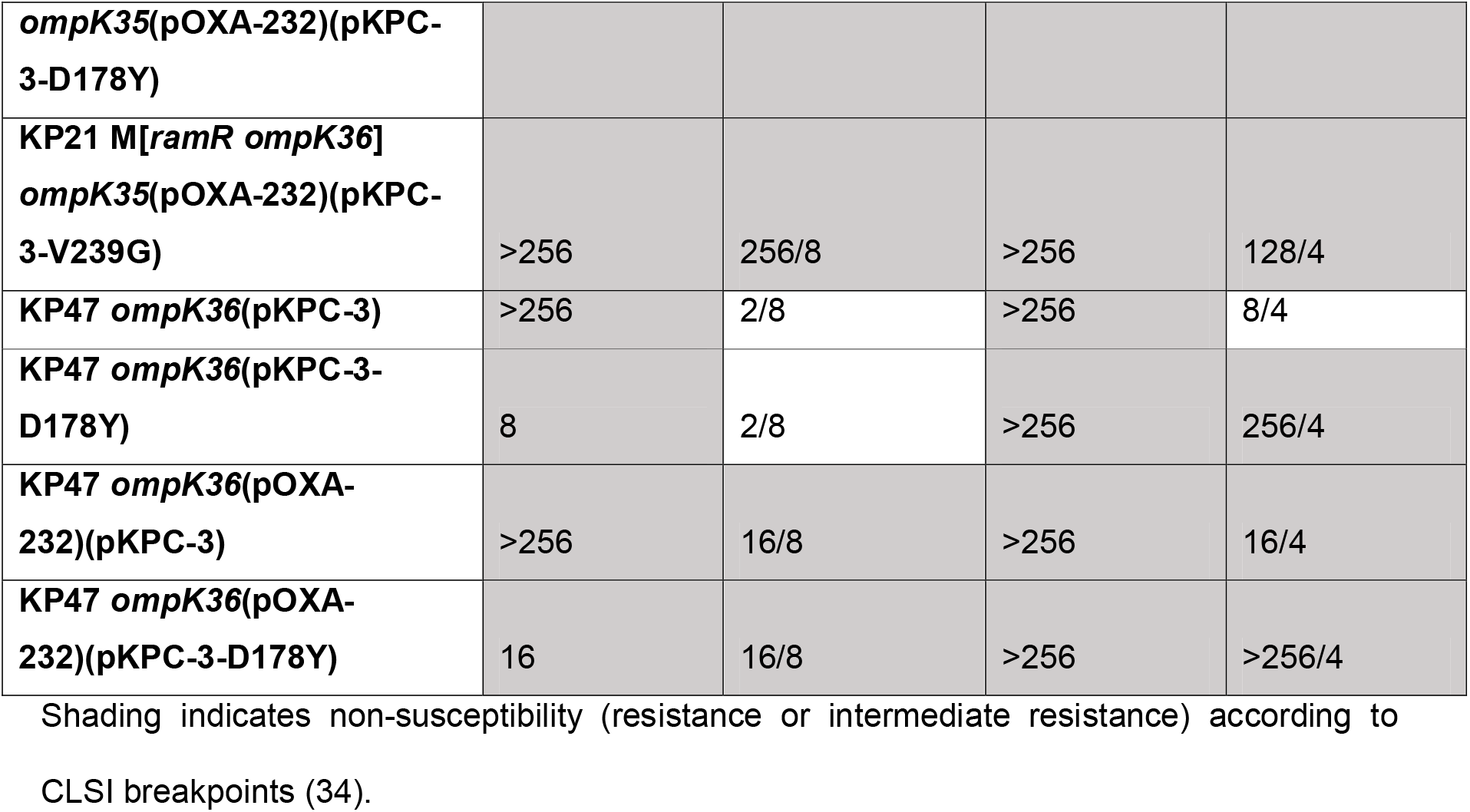
MICs (μg.mL^−1^) of meropenem with or without vaborbactam and of ceftazidime with or without avibactam against derivatives of *K. pneumoniae* clinical isolates.

The meropenem/vaborbactam resistant KP21(pOXA-232) *ompK36* mutant derivatives remained susceptible to ceftazidime (**Table 2**) as expected since OXA-232 is a weak cephalosporinase (26). Also as expected, introduction of a low-copy-number vector carrying *bla*_CTX-M-14_ alongside its native promoter into either KP21(pOXA-232), KP21 *ompK36*(pOXA-232) or KP21 M[*ompK36*](pOXA-232) conferred ceftazidime resistance. The derivatives remained ceftazidime susceptible in the presence of 4 μg.mL^−1^ avibactam (**Table 2**) which is a potent inhibitor of CTX-M (11). Replacement of *bla*_CTX-M-14_ with a gene encoding a variant associated with reduced ceftazidime/avibactam susceptibility, CTX-M-14-P170S (16) drove the ceftazidime MIC against KP21 *ompK36*(pOXA-232) and KP21 M[*ompK36*](pOXA-232) in the presence of avibactam up to 8 μg.mL^−1^, which is still susceptible, but only one doubling dilution below the breakpoint for resistance. Disruption of *ompK35* in KP21 M[*ompK36*](pOXA-232)(pCTX-M-14-P170S) did not further increase the ceftazime/avibactam MIC, confirming that *ramR* loss-of-function phenotypically mimics *ompK35* loss-of-function in this regard (**Table 2**). Accordingly, it was not possible to generate derivatives resistant to both meropenem/vaborbactam and ceftazidime/avibactam using a CTX-M-14 variant associated with reduced susceptibility to ceftazidime/avibactam, even in a *ramR* plus *ompK36* double mutant background co-producing OXA-232.

Introduction of KPC-3 and its derivatives associated with reduced ceftazidime/avibactam susceptibility into KP21(pOXA-232) was also attempted. In this case, both KPC-3 derivatives tested (D178Y or V239G) conferred ceftazidime/avibactam resistance. The D178Y derivative did this at the expense of reducing meropenem MIC into the intermediate-resistant zone. Essentially, there is a trade-off between meropenem hydrolytic activity and reduced avibactam inhibition (or perhaps increased ceftazidime hydrolytic activity) as reported previously (14). The V239G mutant did not suffer from such a large drop in meropenem MIC, but neither S178Y or V239G significantly altered the ability of vaborbactam to inhibit KPC-3 as confirmed because all KP21(pOXA-232) derivatives carrying these KPC-3 variants were meropenem/vaborbactam susceptible (**Table 2**). Accordingly, it was not possible to generate derivatives resistant to both meropenem/vaborbactam and ceftazidime/avibactam using KPC-3 variants associated with ceftazidime/avibactam resistance in a background having wild-type *ompK36*, even with co-production of OXA-232.

### Combining ramR loss, OXA-232, ompK36 loss and KPC-3-D178Y confers dual meropenem/vaborbactam plus ceftazidime/avibactam resistance in K. pneumoniae

The above observations left us with the hypothesis that adding a KPC-3 derivative associated with ceftazidime/avibactam resistance to a background where meropenem non-susceptibility is conferred by mutation of *ramR* (i.e. causing OmpK35 downregulation) *ompK36* loss and OXA-232 production would generate a derivative resistant to ceftazidime/avibactam and meropenem/vaborbactam when used separately. This proved to be correct. MICs of meropenem/vaborbactam and ceftazidime/avibactam were ≥64 μg.mL^−1^ against KP21 M[*ompK36*](pOXA-232) or KP21 *ompK36*(pOXA-232) carrying either the D178Y or V239G derivatives of KPC-3. Additional disruption of *ompK35* only marginally further increased MICs, again confirming that OmpK35 down-regulation caused by *ramR* loss-of-function gives essentially the same phenotype as *ompK35* loss-of-function (**Table 2**).

Despite our use of OXA-232-producing strains for this analysis, production of OXA-232 was not found to be essential for resistance to meropenem/vaborbactam and ceftazidime/avibactam, when used separately. After constructing KP21 *ompK36*(pKPC-3-D178Y) and KP21 *ompK36*(pKPC-3-V239G) – i.e. lacking OXA-232 – we found them to be resistant to both β-lactam/β-lactamase inhibitor pairs (**Table 2**). Another interesting finding from our analysis was that ceftazidime/avibactam and meropenem/vaborbactam resistance can be conferred in the presence of wild-type KPC-3 and OXA-232 in this *ramR* background also lacking *ompK36* (**Table 2**). Clinical *K. pneumoniae* isolates with *ompK35* and *ompK36* loss-of-function mutations and elevated KPC-3 production have recently been identified that are resistant to ceftazidime/avibactam (30) which supports our conclusion that KPC mutations are not essential for resistance.

Finally, we wanted to test whether the two β-lactam/inhibitor pairs would work synergistically, to give hope of clinical efficacy for the double combination therapy that has been discussed as a possibility for clinical use in the literature (20,21). Checkerboard assays (**Figure**) confirmed that KP21 M[*ompK36*](pOXA-232) and KP21 *ompK36*(pOXA-232) carrying pCTX-M14-P170S, pKPC-3 or pKPC-3-V239G are all susceptible to meropenem (MIC ≤ 8 μg.mL^−1^) in the presence of vaborbactam plus ceftazidime/avibactam suggesting that combined therapy would still work. However, KP21 M[*ompK36*](pOXA-232) carrying pKPC-3-D178Y and KP21 *ompK36*(pOXA-232) carrying pKPC-3-D178Y were resistant to meropenem (MIC = 16 μg.mL^−1^) and ceftazidime (MIC ≥ 64 μg.mL^−1^) in the presence of vaborbactam plus avibactam. Further disruption of *ompK35* did not alter this result, again confirming that *ramR*-loss-mediated OmpK35 downregulation (as seen in all KP21 derivatives [29]) mimics the phenotypic effect of *ompK35* loss.

**Figure.**
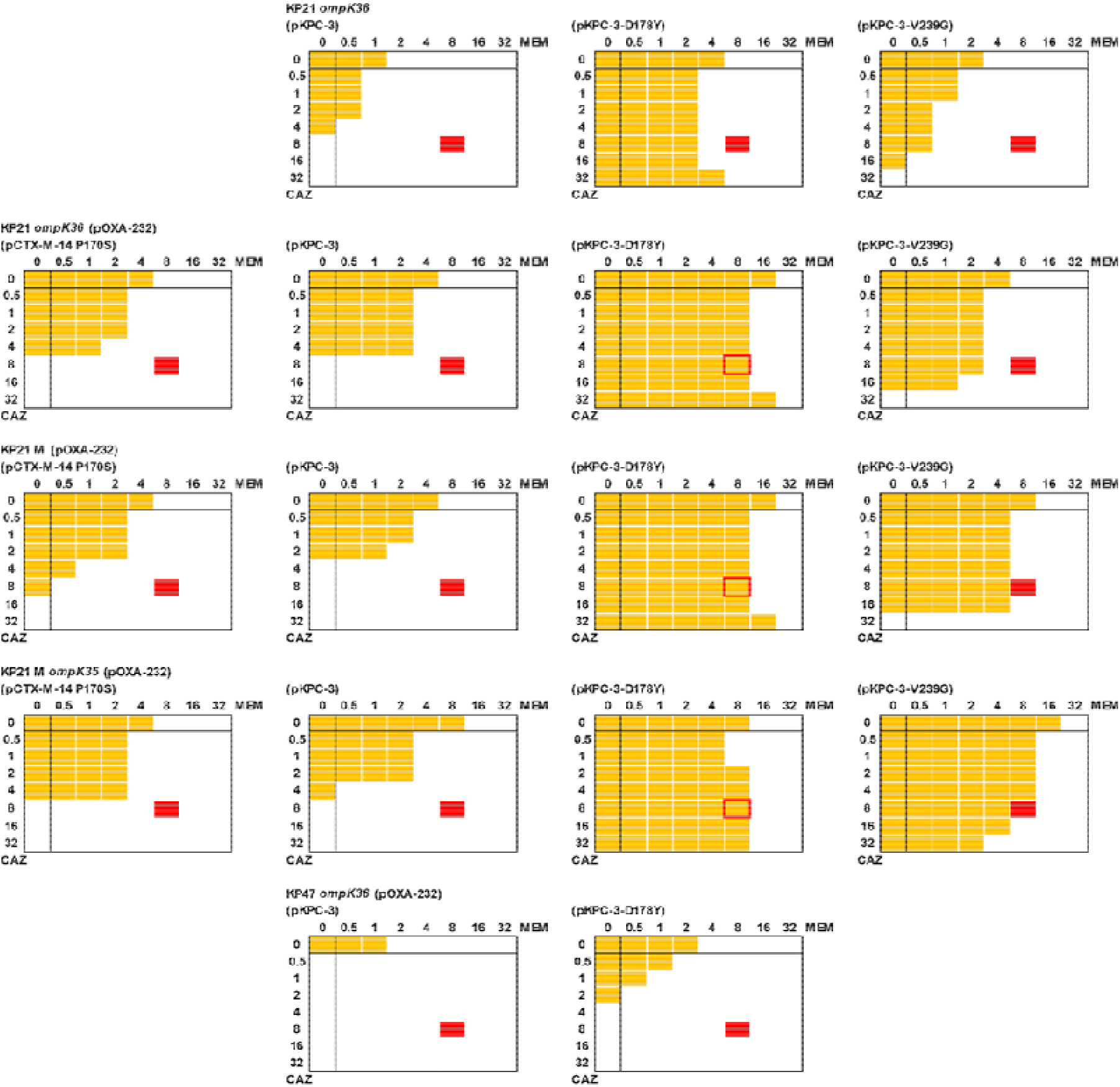
Checkerboard assays for ceftazidime and meropenem in the presence of avibactam and vaborbactam. Each image represents duplicate assays for an 8×8 array of wells in a 96-well plate. All wells contained CA-MHB including avibactam (4 μg.mL^−1^) and vaborbactam (8 μg.mL^−1^). A serial dilution of meropenem (MEM, x-axis) and ceftazidime (CAZ, y-axis) was created from 32 μg.mL^−1^ in each plate as recorded. All wells were inoculated with a suspension of bacteria, made as per CLSI microtiter MIC guidelines (33), and the plate was incubated at 37°C for 20h. Growth was recorded by measuring OD_600_ and growth above background (broth) is recorded as a yellow block. Growth at 8 μg.mL^−1^ ceftazidime and 8 μg.mL^−1^ meropenem (this position indicated in red) in the presence of vaborbactam and avibactam defines resistance based on CLSI breakpoints (34). Bacterial suspensions used were: for images in the top row, KP21[*ramR*] *ompK36*; second row, KP21[*ramR*] *ompK36*(pOXA-232); third row, KP21[*ramR*] M[*ompK36*](pOXA-232); fourth row, KP21[*ramR*] M[*ompK36*] *ompK35*(pOXA-232); fifth row, KP47 *ompK36*(pOXA-232). In each case, bacteria also carry the following plasmids (where tested): images in first column, pCTX-M-14 P170S; second column, pKPC-3; third column, pKPC-3-D178Y; fourth column, pKPC-3-V239G.

We have, therefore, generated *K. pneumoniae* derivatives resistant to ceftazidime/avibactam plus meropenem/vaborbactam when both pairs are used together. This was achieved in four steps relative to wild-type (*ramR* loss-of-function, acquisition of OXA-232, *ompK36* loss-of-function, acquisition of KPC-3-D178Y) each replicating an event previously identified in clinical *K. pneumoniae* isolates. When we constructed a derivative of clinical isolate KP47 (wild type for *ramR*) carrying OXA-232 and KPC-3-D178Y and with *ompK36* inactivated, the derivative was meropenem/vaborbactam and ceftazidime/avibactam resistant when the pairs were used separately (**Table 2**) but not when both pairs were used at the same time (**Figure**). This confirms that OmpK35 downregulation caused by *ramR* loss-of-function is necessary to confer this dual-resistant phenotype. When we constructed the derivative KP21 *ompK36* carrying KPC-3-D178Y or KPC-3-V239G but without carriage of OXA-232 it was also resistant to both pairs when used separately (**Table 2**) but not resistant when the pairs were used together (**Figure**). Therefore, production of an OXA-48 like enzyme is also necessary for dual resistance.

Overall, this work confirms the remarkable capacity of *K. pneumoniae* to acquire resistance to the latest combination therapies available in the clinic by layering resistance mechanisms. Essentially, one low frequency step (specific mutations in a gene) is needed for ceftazidime/avibactam resistance; three steps (two high frequency loss-of-function mutations and a gene acquisition event, who’s frequency depends on many factors) are needed for meropenem/vaborbactam resistance. If one of the three steps is acquisition of a KPC associated with ceftazidime/avibactam resistance, then these three steps lead to resistance to both ceftazidime/avibactam and meropenem/vaborbactam when used separately; finally adding in an fourth step: acquisition of an OXA-48-like carbapenemase is all that is needed to give resistance to both β-lactam/β-lactamase inhibitor pairs when used together. Of course, the order that these steps occur does not affect the result. So, it would seem prudent to make every effort to identify, via molecular diagnostics, intermediate stages in this acquisition process, e.g. *ramR* or *ompK35* mutants carrying KPC or an OXA-48-like enzyme. It would also seem prudent not to rely on sequential use of β-lactam/β-lactamase inhibitor pairs, which might select for the dual-resistant phenotype observed here. In **Table 3** we attempt to address the issue of how different starting genotypes influence the chances of obtaining derivatives with specific resistance phenotypes. Whilst the frequencies reported for plasmid transmission events will vary widely dependent on local plasmid ecology, we hope that this table stimulates discussion around clinical decision making to reduce the emergence of resistance at the site of infection or in the patient’s gut flora. And will increase the desire to obtain relevant genotypic information about circulating bacterial populations, and even the ecology of resistant organisms in the patient’s gut microbiome prior to choosing therapy.

**Table 3.** Rough relative estimates for risk of acquiring resistance to β-lactam/β-lactamase inhibitor pairs, singularly or in combination based on starting genotype.

| Phenotype > | MER/VAB | AVI/CAZ | MER/VAB and AVI/CAZ separately | MER/VAB and AVI/CAZ together |
| --- | --- | --- | --- | --- |
| **Genotype Needed >** | *ramR/ompK35+ompK36+KPC/OXA-232* | KPC-MUT **or**<br>*ramR/ompK35+ompK36+KPC+OXA-232* | *ramR/ompK35+ompK36+KPC+OXA-232* **or** *ramR/ompK35+ompK36+KPC-MUT* | *ramR/ompK35+ompK36+KPC-MUT+OXA-232* |
| **Actual Genotype** | **Needed (est. frequency)** | **Needed (est. frequency)** | **Needed (est. frequency)** | **Needed (est. frequency)** |
| WT+KPC | *ramR/ompK35+OmpK36* ($10^{-14}$) | KPC-MUT ($10^{-9}$) | *ramR/ompK35+ompK36+KPC-MUT* ($10^{-23}$) | *ramR/ompK35+ompK36+KPC-MUT+OXA-232* ($10^{-33}$)** |
| | | *ramR/ompK35+ompK36+OXA-232* ($10^{-23}$)** | *ramR/ompK35+ompK36+OXA-232* ($10^{-23}$)** | *ramR+ompK36+KPC-MUT+OXA-232* ($10^{-33}$)** |
| *ramR/ompK35+KPC* | *ompK36* ($10^{-7}$) | KPC-MUT ($10^{-9}$) | *ompK36+KPC-MUT* ($10^{-16}$) | *ompK36+KPC-MUT+OXA-232* ($10^{-26}$)** |
| | | *ompK36+OXA-232* ($10^{-16}$)** | *ompK36+OXA-232* ($10^{-16}$)** | *ompK36+KPC-MUT+OXA-232* ($10^{-26}$)** |
| *ompK36+KPC* | *ramR/ompK35* ($10^{-7}$) | KPC-MUT ($10^{-9}$) | *ramR/ompK35+KPC-MUT* ($10^{-16}$) | *ramR/ompK35+KPC-MUT+OXA-232* ($10^{-26}$)** |
| | | *ramR/ompK35+OXA-232* ($10^{-16}$)** | *ramR/ompK35+OXA-232* ($10^{-16}$)** | *ompK36+KPC-MUT+OXA-232* ($10^{-26}$)** |
| WT+KPC-MUT | *ramR/ompK35+OmpK36* ($10^{-14}$) | -- | *ramR/ompK35+OmpK36* ($10^{-14}$) | *ramR/ompK35+OmpK36+OXA-232* ($10^{-24}$)** |
| *ramR/ompK35+KPC-MUT* | *ompK36* ($10^{-7}$) | -- | *ompK36* ($10^{-7}$) | *ompK36+OXA-232* ($10^{-17}$)** |
| *ompK36+KPC-MUT* | *ramR/ompK35* ($10^{-7}$) | -- | *ramR/ompK35* ($10^{-7}$) | *ramR/ompK35+OXA-232* ($10^{-17}$)** |
| *ramR/ompK35+ompK36+KPC* | -- | KPC-MUT ($10^{-9}$) | KPC-MUT ($10^{-9}$) | KPC-MUT+OXA-232 ($10^{-19}$)** |
| | | OXA-232 ($10^{-10}$)** | OXA-232 ($10^{-10}$)** | KPC-MUT+OXA-232 ($10^{-20}$)** |
| WT+KPC+OXA-232 | *ramR/ompK35+OmpK36* ($10^{-14}$) | KPC-MUT ($10^{-9}$) | *ramR/ompK35+OmpK36+KPC-MUT* ($10^{-23}$) | *ramR/ompK35+OmpK36+KPC-MUT* ($10^{-23}$) |
| | | *ramR/ompK35+OmpK36* ($10^{-14}$) | *ramR/ompK35+OmpK36* ($10^{-14}$) | *ramR/ompK35+OmpK36+KPC-MUT* ($10^{-23}$) |
| *ramR/ompK35+* | *ompK36* ($10^{-7}$) | KPC-MUT ($10^{-9}$) | *ompK36+KPC-MUT* ($10^{-16}$) | *ompK36+KPC-MUT* ($10^{-16}$) |
| KPC+OXA-232 |  |  |  |  |
| | | *ompK36* ($10^{-7}$) | *ompK36* ($10^{-7}$) | *ompK36*+KPC-MUT ($10^{-16}$) |
| *ompK36*+KPC+OXA-232 | *ramR/ompK35* ($10^{-7}$) | KPC-MUT ($10^{-9}$) | *ramR/ompK35*+KPC-MUT ($10^{-16}$) | *ramR/ompK35*+KPC-MUT ($10^{-16}$) |
| | | *ramR/ompK35* ($10^{-7}$) | *ramR/ompK35* ($10^{-7}$) | *ramR/ompK35*+KPC-MUT ($10^{-16}$) |
| WT+KPC-MUT+OXA-232 | *ramR/ompK35+OmpK36* ($10^{-14}$) | -- | *ramR/ompK35+OmpK36* ($10^{-14}$) | *ramR/ompK35+OmpK36* ($10^{-14}$) |
| *ramR/ompK35*+KPC-MUT+OXA-232 | *ompK36* ($10^{-7}$) | -- | *ompK36* ($10^{-7}$) | *ompK36* ($10^{-7}$) |
| *ompK36*+KPC-MUT+OXA-232 | *ramR/ompK35* ($10^{-7}$) | -- | *ramR/ompK35* ($10^{-7}$) | *ramR/ompK35* ($10^{-7}$) |

\*\*The risk of plasmid transfer is impossible to assess. It will depend, amongst many other factors, on the transmissibility and abundance of the plasmid that must be transferred. KPC plasmids of the pKpQIL type, in perfect laboratory conditions transfer at a frequency of $10^{-6}$ to $10^{-8}$ transconjugants per donor cell (40); IncL/M OXA-48 plasmids transfer is at a frequency of almost $10^{-6}$ (41). So, in mixed populations, such as might be seen in the gut microbiota, we have assessed the maximum possible transfer frequency as $10^{-10}$ for each. Of course, if a patient is clear of any plasmid-encoded KPC or OXA-48 producing bacteria, the observed transfer frequency for the genes encoding these enzymes will be close to zero. Frequency of spontaneous resistance in KPC was equated with the observed mutation frequency ( $10^{-9}$ ) for streptomycin resistance in *Escherichia coli*, since this represents a similarly specific set of mutations (42). A mutator phenotype could increase this by two orders of magnitude (43). We set the frequency of loss-of-function mutation (*ramR*, *ompK35* or *ompK36*) as 100-fold higher than the spontaneous mutation frequency because of the number of possible mutations that could disrupt a gene, but this is likely to be a very conservative estimate as previously described (44). Abbreviations: MER, meropenem, VAB, vaborbactam, CAZ, ceftazidime, AVI, avibactam, KPC-MUT, KPC derivatives associated with AVI/CAZ resistance.

## Experimental

### Materials, bacterial isolates and plasmids

Chemicals were from Sigma and growth media from Oxoid, unless otherwise stated. Meropenem was from Sequoia Research Products, vaborbactam and avibactam were from MedChemExpress. Strains used were *K. pneumoniae* Ecl8 (31) the TEM-1-producing *ramR* mutant (Arg44FS) clinical isolate KP21, the wild-type clinical isolate KP47 and the in vitro-selected Ecl8-derived *oqxR* (Tyr109STOP) or *ramR* (Thr124Pro) loss-of-function mutants (29). A plasmid carrying *bla*_*OXA*-232_ (pOXA-232) was recovered from *K. pneumoniae* clinical isolate KP11 (29) using a Qiagen plasmid purification kit, with the plasmid then used to transform *E. coli* DH5α to reduced piperacillin/tazobactam susceptibility (8 μg.mL^−1^ piperacillin and 4 μg.mL^−1^ tazobactam) using electroporation. Recombinants were confirmed to be cefotaxime susceptible before the plasmid was re-purified and used to transform *K. pneumoniae* isolates. The *bla*_*OXA*-232_-encoding region was confirmed as being unchanged from the original using PCR sequencing.

### Selection and generation of mutants

To select meropenem/vaborbactam resistant mutants, 100 μL aliquots of overnight cultures of KP21 grown in Nutrient Broth (NB) were spread onto Mueller Hinton agar containing 16 μg.mL^−1^ meropenem in the presence of 8 μg.mL^−1^ of vaborbactam, which were then incubated for 24 h. Insertional inactivation of *ompK35* or *ompK36* was performed using the pKNOCK suicide plasmid (32). The *ompK35* and *ompK36* DNA fragments were amplified with Phusion High-Fidelity DNA Polymerase (NEB, UK) from *K. pneumoniae* Ecl8 genomic DNA using primers *ompK35* KO FW (5′-TCCCAGACCACAAAAACCCG-3′) and *ompK35* KO RV (5′-CCAGACCGAAGAAGTCGGAG-3′); *ompK36* KO FW (5′-CGTTCAGGCGAACAACACTG-3′) and *ompK36* KO RV (5′-AAGTTCAGGCCGTCAACCAG-3′). Each PCR product was ligated into the pKNOCK-GM at the SmaI site. The recombinant plasmid was then transferred into wild-type *K. pneumoniae* isolates by conjugation. Mutants were selected for gentamicin non-susceptibility (5 μg.mL^−1^) and the mutation was confirmed by PCR using primers *ompK35* full length FW (5′-CACTTCGATGTATTTAACCAG-3′) and *ompK35* full length RV (5′-ATGATGAAGCGCAATATTCTG-3′) *ompK36* full length FW (5′-GAGGCATCCGGTTGAAATAG-3′) and *ompK36* full length RV (5′-ATTAATCGAGGCTCCTCTTAC-3′).

### Determining MICs of antimicrobials and checkerboard assays

MICs were determined using CLSI broth microtiter assays (33) and interpreted using published breakpoints (34). Checkerboard assays were performed using an adapted microtiter MIC assay. Briefly, a PBS bacterial suspension was prepared to obtain a stock of OD_600_=0.01. The final volume in each well of a 96-well cell culture plate (Corning Costar) was 200 μL and included 20 μL of the bacterial suspension. All wells contained Cation Adjusted Muller Hinton broth (CA-MHB) with avibactam (4 μg.mL^−1^) and vaborbactam (8μg.mL^−1^) with a serial dilution of meropenem from right to left and ceftazidime from the bottom to the top of the plate. Bacterial growth was determined after 20 h of incubation by measuring optical density at 600 nm (OD_600_) using a POLARstar Omega spectrophotometer (BMG Labtech).

### Proteomics

Five hundred microlitres of an overnight CA-MHB culture were transferred to 50 mL CA-MHB and cells were grown at 37°C to 0.6 OD_600_. Cells were pelleted by centrifugation (10 min, 4,000 × *g*, 4°C) and resuspended in 20 mL of 30 mM Tris-HCl, pH 8 and broken by sonication using a cycle of 1 s on, 0.5 s off for 3 min at an amplitude of 63% using a Sonics Vibracell VC-505TM (Sonics and Materials Inc., Newton, Connecticut, USA). The sonicated samples were centrifuged at 8,000 rpm (Sorval RC5B PLUS using an SS-34 rotor) for 15 min at 4°C to pellet intact cells and large cell debris. For envelope preparations, the supernatant was subjected to centrifugation at 20,000 rpm for 60 min at 4°C using the above rotor to pellet total envelopes. To isolate total envelope proteins, this total envelope pellet was solubilized using 200 μL of 30 mM Tris-HCl, pH 8 containing 0.5% (w/v) SDS.

Protein concentrations in all samples were quantified using Biorad Protein Assay Dye Reagent Concentrate according to the manufacturer’s instructions. Proteins (5 μg/lane for envelope protein analysis) were separated by SDS-PAGE using 11% acrylamide, 0.5% bis-acrylamide (Biorad) gels and a Biorad Min-Protein Tetracell chamber model 3000X1. Gels were resolved at 200 V until the dye front moved approximately 1 cm into the separating gel. Proteins in all gels were stained with Instant Blue (Expedeon) for 20 min and de-stained in water.

The 1 cm of gel lane was subjected to in-gel tryptic digestion using a DigestPro automated digestion unit (Intavis Ltd). The resulting peptides from each gel fragment were fractionated separately using an Ultimate 3000 nanoHPLC system in line with an LTQ-Orbitrap Velos mass spectrometer (Thermo Scientific). In brief, peptides in 1% (v/v) formic acid were injected onto an Acclaim PepMap C18 nano-trap column (Thermo Scientific). After washing with 0.5% (v/v) acetonitrile plus 0.1% (v/v) formic acid, peptides were resolved on a 250 mm × 75 μm Acclaim PepMap C18 reverse phase analytical column (Thermo Scientific) over a 150 min organic gradient using 7 gradient segments (1-6% solvent B over 1 min, 6-15% B over 58 min, 15-32% B over 58 min, 32-40% B over 5 min, 40-90% B over 1 min, held at 90% B for 6 min and then reduced to 1% B over 1 min) with a flow rate of 300 nL/min. Solvent A was 0.1% formic acid and Solvent B was aqueous 80% acetonitrile in 0.1% formic acid. Peptides were ionized by nano-electrospray ionization MS at 2.1 kV using a stainless-steel emitter with an internal diameter of 30 μm (Thermo Scientific) and a capillary temperature of 250°C. Tandem mass spectra were acquired using an LTQ-Orbitrap Velos mass spectrometer controlled by Xcalibur 2.1 software (Thermo Scientific) and operated in data-dependent acquisition mode. The Orbitrap was set to analyse the survey scans at 60,000 resolution (at m/z 400) in the mass range m/z 300 to 2000 and the top twenty multiply charged ions in each duty cycle selected for MS/MS in the LTQ linear ion trap. Charge state filtering, where unassigned precursor ions were not selected for fragmentation, and dynamic exclusion (repeat count 1; repeat duration 30 s; exclusion list size 500) were used. Fragmentation conditions in the LTQ were as follows: normalized collision energy of 40%; activation q 0.25; activation time 10 ms; and minimum ion selection intensity 500 counts.

The raw data files were processed and quantified using Proteome Discoverer software v1.4 (Thermo Scientific) and searched against the UniProt *K. pneumoniae* strain ATCC 700721 / MGH 78578 database (5126 protein entries; UniProt accession 272620) using the SEQUEST (Ver. 28 Rev. 13) algorithm. Peptide precursor mass tolerance was set to 10 ppm, and MS/MS tolerance was set to 0.8 Da. Search criteria included carbamidomethylation of cysteine (+57.0214) as a fixed modification and oxidation of methionine (+15.9949) as a variable modification. Searches were performed with full tryptic digestion and a maximum of 1 missed cleavage was allowed. The reverse database search option was enabled and all peptide data was filtered to satisfy false discovery rate (FDR) of 5%. The Proteome Discoverer software generates a reverse “decoy” database from the same protein database used for the analysis and any peptides passing the initial filtering parameters that were derived from this decoy database are defined as false positive identifications. The minimum cross-correlation factor filter was readjusted for each individual charge state separately to optimally meet the predetermined target FDR of 5 % based on the number of random false positive matches from the reverse decoy database. Thus, each data set has its own passing parameters. Protein abundance measurements were calculated from peptide peak areas using the Top 3 method (35) and proteins with fewer than three peptides identified were excluded. The proteomic analysis was repeated three times for each parent and mutant strain, each using a separate batch of cells.

### Whole genome sequencing to Identify mutations

Whole genome resequencing was performed by MicrobesNG (Birmingham, UK) on a HiSeq 2500 instrument (Illumina, San Diego, CA, USA). Reads were trimmed using Trimmomatic (36) and assembled into contigs using SPAdes 3.10.1 (http://cab.spbu.ru/software/spades/). Assembled contigs were mapped to the *K. pneumoniae* Ecl8 reference genome (GenBank accession number GCF_000315385.1) by using progressive Mauve alignment software (37).

### Cloning bla_CTX-M-14_ and bla_KPC-3_ and site-directed mutagenesis

*bla*_CTX-M-14_ was amplified from a human urinary *E. coli* isolated from primary care (38) by PCR with Phusion High-Fidelity DNA Polymerase (NEB, UK) using primers CTX-M-14 FW (5′-CCGGAATTCAATACTACCTTGCTTTCTGA-3′) and CTX-M-14 RV (5′-CCGGAATTCCGTAGCGGAACGTTCATCAG-3′) and ligated into pUBYT (39) at the EcoRI site. CTX-M-14 site-directed mutagenesis (*bla*_CTX-M-14-P170S_) was performed using the QuikChange Lightning Site-Directed Mutagenesis Kit (Agilent, USA) with the primer CTX-M-14-P170S-FW (5′-TCTGGATCGCACTGAATCTACGCTGAATACCGC-3′).

*bla*_KPC-3_ was amplified from pKpQIL isolated from *K. pneumoniae* KP30 (29) as above using primers KPC-3 FW (5′-CCGGAATTCGTAAAGTGGGTCAGTTTTCAG-3′) and KPC-3 RV (5′-GGCTCTGAAAATCATCTATTGGAATTCCGG-3′) and ligated into pUBYT at the EcoRI site. KPC-3 site-directed mutagenesis (*bla*_KPC-3-D178Y_ and *bla*_KPC-3-V239G_) was performed using a two-step, PCR-based site-directed mutagenesis strategy. *bla*_KPC-3-D178Y_ was constructed using primers KPC-3-D178Y-FW (5′-AGGCGATGCGCGCTATACCTCATCGCC-3′) and KPC-3-D178Y-RV (5′-GGCGATGAGGTATAGCGCGCATCGCCT-3′); *bla*_KPC-3-V239G_ was constructed using primers KPC-3-V239G-FW (5′-GGAACCTGCGGAGGGTATGGCACGGCA-3′) and KPC-3-V239G-RV (5′-TGCCGTGCCATACCCTCCGCAGGTTCC-3′). Carriage of all pUBYT plasmids was selected using kanamycin (30 μg.mL^−1^)

## Acknowledgments

This work was funded by grant MR/S004769/1 to M.B.A. from the Antimicrobial Resistance Cross Council Initiative supported by the seven United Kingdom research councils and the National Institute for Health Research. Additional funding came via a bequest from the estate of the late Professor Graham Ayliffe. Genome sequencing was provided by MicrobesNG (http://www.microbesng.uk/), which is supported by the BBSRC (grant number BB/L024209/1). We are grateful to Dr Kate Heesom, School of Biochemistry, University of Bristol, for performing the proteomics analysis and to Dr Jacqueline Findlay, School of Cellular & Molecular Medicine, for purifying pOXA-232.

**We declare no conflicts of interest**

